# Reformulating Protein Corona Prediction as Universal Protein-conditioned Learning

**DOI:** 10.64898/2026.10.09.752255

**Authors:** Xinhe Wu, Chen Wang, Yadong Jin, Yuliang Zhao, Huan Tang

## Abstract

When nanoparticles enter biological environments, their surfaces are rapidly coated by proteins, forming a protein corona that governs their biological identity and downstream responses. Existing computational methods treat each protein as an independent prediction task, preventing the learning of shared adsorption principles and limiting generalization beyond experimentally characterized proteins. Here we introduce NanoPCT, a universal protein-conditioned learning framework that jointly models nanoparticle physicochemical properties and protein sequence representations. Using frozen ESM-2 embeddings and a joint interaction transformer, NanoPCT learns transferable representations of nano–bio interaction patterns across hundreds of proteins and maintains robust performance as protein diversity increases. The learned protein representation further enables zero-shot prediction for previously unseen proteins and biologically distinct protein groups through classifier reconstruction in embedding space. Beyond prediction, NanoPCT supports large-scale in silico inverse design and reveals consistent adsorption preferences between observed and unseen protein populations, suggesting that it captures transferable interaction patterns rather than protein-specific memorization. NanoPCT provides a unified framework for protein corona prediction, zero-shot generalization, and rational nanomaterial design.

## 1 Introduction

Engineered nanomaterials have emerged as a central platform for drug delivery, molecular imaging, diagnostics, and immunotherapy, offering unprecedented opportunities for precision medicine [1–3]. Yet despite remarkable advances in nanoparticle engineering, translating nanomaterials from *in vitro* optimization to *in vivo* efficacy remains notoriously difficult [4–6]. A major reason is that nanoparticles rarely interact with biological systems in their synthetic form. Instead, immediately upon exposure to physiological fluids, their surfaces become coated by hundreds of biomolecules, predominantly proteins, forming the so-called protein corona [7, 8]. This dynamic biological interface effectively replaces the original physicochemical identity of the nanoparticle and governs its cellular uptake, biodistribution, immune recognition, circulation lifetime, and ultimately therapeutic efficacy [9, 10]. Consequently, the ability to predict protein corona formation has become one of the central challenges in nanomedicine, as it determines how engineered nanomaterials are ultimately perceived by living systems [11].

Experimental characterization of protein corona composition, typically through liquid chromatography coupled with tandem mass spectrometry (LC-MS/MS), has substantially advanced our understanding of nano-bio interactions [12, 13]. However, exhaustive experimental profiling remains prohibitively expensive and inconsistent owing to the enormous combinatorial space generated by nanoparticle physicochemical properties, protein diversity, and biological environments [14, 15]. Machine learning has therefore emerged as an attractive strategy for accelerating protein corona prediction [16, 17]. Nevertheless, nearly all existing computational approaches formulate protein corona prediction as a collection of independent protein-specific tasks, training a separate predictor for each target protein [11, 16, 18–22]. Although effective for proteins represented in the training data, this formulation implicitly treats every protein as an isolated prediction problem, preventing knowledge sharing across proteins and fundamentally limiting prediction to proteins and biological contexts represented in existing datasets. As a consequence, existing models are unable to generalize to the vast majority of proteins for which no adsorption measurements are available. This limitation is particularly consequential because the experimentally characterized protein corona represents only a small fraction of the biological protein space. In practical nanomedicine applications, nanoparticles encounter highly diverse and context-dependent protein environments, including proteins that have never been experimentally profiled against engineered nanomaterials. Therefore, a clinically relevant predictive framework must not only interpolate among previously characterized proteins, but also infer adsorption behavior for entirely unseen proteins without requiring additional experimental measurements. We argue that this limitation originates not from model capacity, but from the formulation of the problem itself. Protein adsorption is governed by shared physicochemical principles arising from nanoparticle–protein interactions at the interface, rather than by unrelated protein-specific rules. Therefore, protein corona prediction should not be regarded as hundreds of independent classification tasks, but instead as a single protein-conditioned learning problem, in which proteins act as continuous descriptors of interaction partners rather than discrete prediction targets. Such a reformulation would allow adsorption knowledge learned from experimentally characterized proteins to be transferred to previously unseen proteins, transforming protein corona prediction from a closed-set recognition problem into an open-world prediction framework where models can estimate interactions across the unexplored protein landscape.

Recent advances in protein language models (PLMs) [23–25] provide the missing representation required to realize this paradigm. Large-scale models such as ESM-2, trained on millions of protein sequences, encode proteins into continuous embedding spaces that preserve evolutionary, structural, and biochemical relationships [23]. Within these learned manifolds, proteins with similar molecular properties occupy neighboring regions, enabling relationships between proteins to be represented geometrically rather than categorically [26, 27]. This continuous representation raises the possibility that adsorption behaviors can also be learned as continuous functions over protein sequence space, allowing predictive knowledge to extend beyond proteins explicitly observed during training. Consequently, protein language models provide a potential solution to one of the central barriers in protein corona prediction: the ability to extrapolate beyond proteins represented in existing experimental datasets. Such a capability would fundamentally redefine protein corona prediction from a closed-set learning problem into an open-world prediction framework.

Here we present NanoPCT (Nanoparticle–Protein Corona Transformer), a universal protein-conditioned learning framework that reformulates protein corona prediction within this new paradigm. Rather than constructing separate models for individual proteins, NanoPCT jointly learns adsorption behaviors for all proteins in a unified architecture by integrating nanoparticle physicochemical descriptors with protein sequence representations derived from a pretrained protein language model. This unified formulation naturally supports zero-shot [28] transfer across both previously unmeasured proteins and biologically distinct protein groups through continuous protein representations, without requiring additional experimental measurements or model retraining.

Using a large-scale protein corona dataset spanning hundreds of proteins and diverse nanomaterials, We demonstrate that universal protein-conditioned learning not only achieves state-of-the-art predictive performance for observed proteins, but also transfers adsorption prediction to previously unseen proteins and functional protein groups. Furthermore, through large-scale *in silico* inverse nanoparticle design, we show that the learned representations reveal transferable material design principles that remain highly consistent between trained and unseen proteins. Together, these results establish a new computational paradigm for protein corona prediction, shifting the field from protein-specific memorization toward transferable representation learning that enables prediction across unexplored protein landscapes and supports rational nanomaterial design beyond existing experimental knowledge.

## 2 Results

### 2.1 Universal protein-conditioned learning

Traditional protein corona prediction methods formulate each protein as an independent prediction task, requiring one model for each target protein [11, 16, 18–22]. Such a formulation prevents knowledge sharing across proteins and makes it impossible to generalize to unseen proteins. We therefore first investigate whether protein corona prediction can instead be formulated as a universal protein-conditioned learning problem, where a single model is trained to predict adsorption for all proteins simultaneously. This formulation is conceptually related to conditional learning and multi-task learning [29], but differs fundamentally in that proteins are represented as continuous biological entities rather than discrete task identities.

#### 2.1.1 Feature-based universal learning

To evaluate whether a universal predictor benefits from increasing protein diversity, we randomly selected different numbers of training proteins (20, 40, 80, 120 and 178) and trained three universal models, including LightGBM [30], XGBoost [31] and NanoPCT. All models were evaluated under the same out-of-fold protocol.

As shown in Fig. 1, all three universal models maintained competitive prediction performance while simultaneously learning an increasing number of proteins. Interestingly, prediction accuracy did not deteriorate as more proteins were incorporated into training. Instead, NanoPCT exhibited remarkably stable performance across all training scales, while the two tree-based models showed only moderate degradation.

**Fig. 1:**
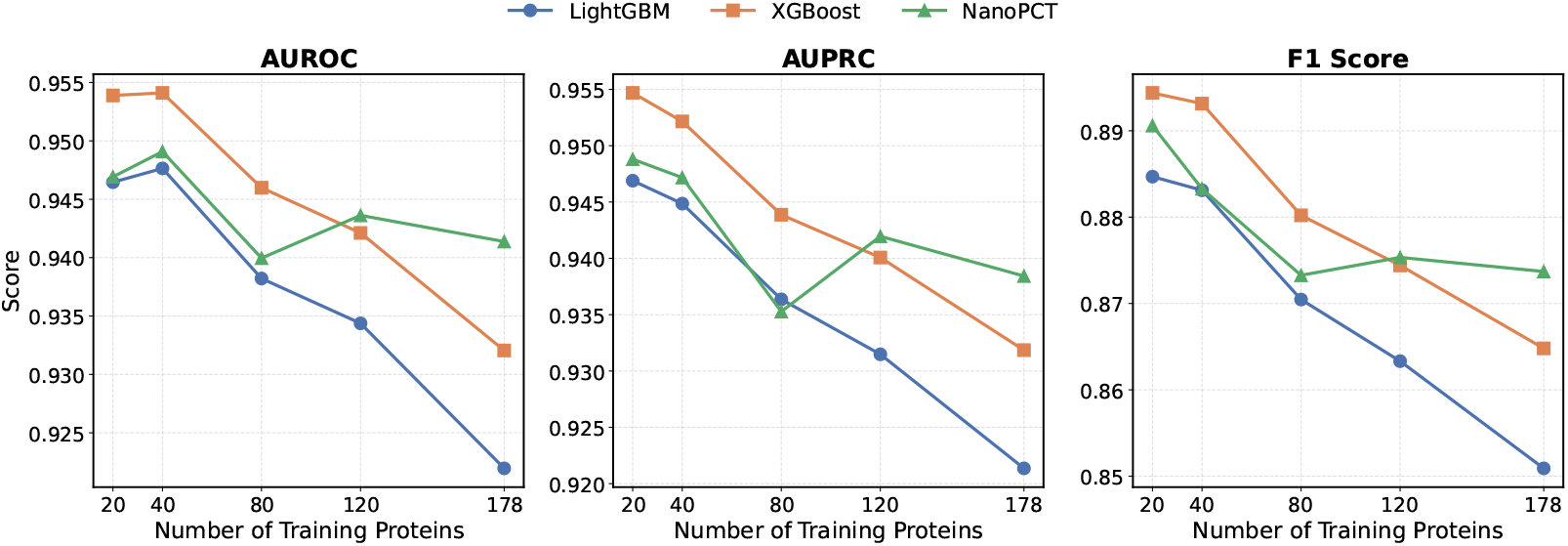
Performance of universal protein-conditioned models as the number of training proteins increases. AUROC, AUPRC and F1-score are reported for LightGBM, XGBoost and NanoPCT.

When trained using all 178 proteins, NanoPCT achieved the highest overall performance, obtaining an AUROC of 0.941, an AUPRC of 0.938 and an F1-score of 0.874. In contrast, LightGBM showed a larger decrease as the number of proteins increased, suggesting that feature-engineered models have more limited capacity for modeling diverse protein adsorption behaviors within a unified predictor.

These results demonstrate that protein corona prediction can be reformulated as a universal learning problem without sacrificing predictive accuracy. More importantly, increasing protein diversity does not introduce severe negative transfer, indicating that adsorption behaviors of different proteins share transferable representations that can be jointly learned by a single model.

#### 2.1.2 Sequence space provides the basis for generalization

The success of universal learning relies on the existence of a continuous representation space where proteins with similar biochemical properties are located close to each other [23–25]. To investigate whether protein language model embeddings satisfy this property, we projected the ESM embeddings of all 393 proteins into two dimensions using both UMAP [32] and t-SNE [33]. We further quantified the relationship between unseen proteins and the training set by measuring the cosine similarity between each unseen protein and its nearest training protein in the embedding space.

As illustrated in Fig. 2a, proteins used for training and proteins reserved for zero-shot evaluation occupy highly overlapping regions in the learned sequence embedding space. Instead of forming two isolated clusters, unseen proteins are consistently intermixed with training proteins across multiple local neighborhoods.

**Fig. 2:**
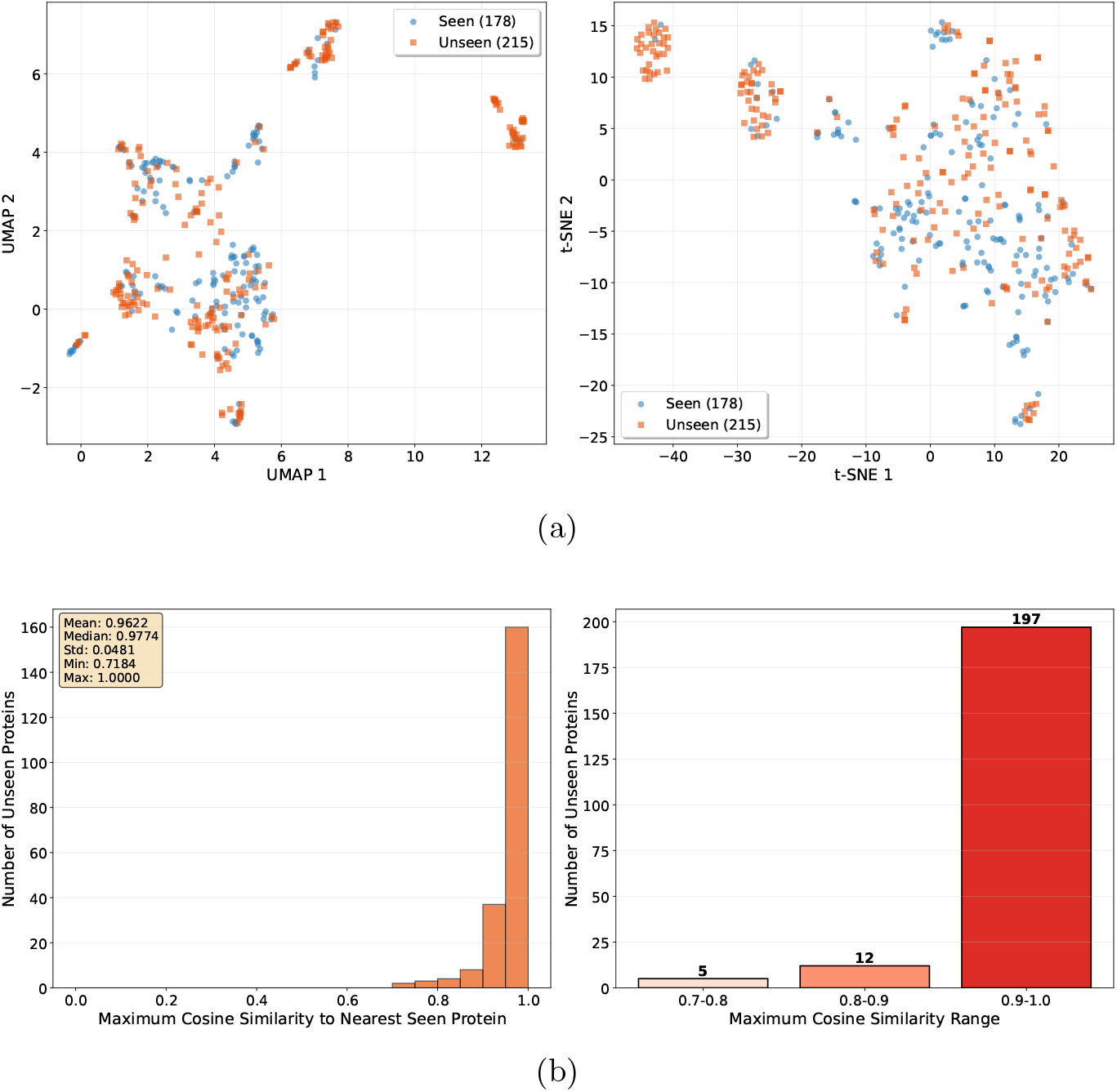
Analysis of the ESM protein embedding space. **(a)**, UMAP projection of protein embeddings, and t-SNE projection of protein embeddings. Blue circles denote proteins used during training, whereas orange squares denote unseen proteins reserved exclusively for zero-shot evaluation. **(b)**, Distribution of the maximum cosine similarity between each unseen protein and its nearest training protein in the embedding space.

This observation suggests that ESM embeddings capture continuous biochemical relationships rather than discrete protein identities. Consequently, unseen proteins naturally reside inside the manifold established by training proteins, providing the geometric foundation for zero-shot generalization.

To quantitatively assess the proximity between unseen and training proteins, we computed the maximum cosine similarity between every unseen protein and its nearest training protein in the embedding space. As shown in Fig. 2b, the nearest-neighbor similarity is consistently high for most unseen proteins. The average maximum cosine similarity reaches 0.962, with 197 out of 215 unseen proteins exhibiting a similarity above 0.9 to at least one training protein. Only five proteins fall below a similarity of 0.8.

These results indicate that the majority of unseen proteins are well represented within the learned sequence space, providing a favorable basis for transferring adsorption knowledge from known proteins to novel proteins. Rather than extrapolating far outside the training distribution, NanoPCT primarily performs local interpolation within a continuous protein manifold, which helps explain its strong zero-shot prediction capability.

### 2.2 NanoPCT architecture

To enable universal protein-conditioned prediction, we developed NanoPCT, a transformer-based framework [34] that jointly models nanoparticle physicochemical properties and protein sequence representations within a unified architecture (Fig. 3).

**Fig. 3:**
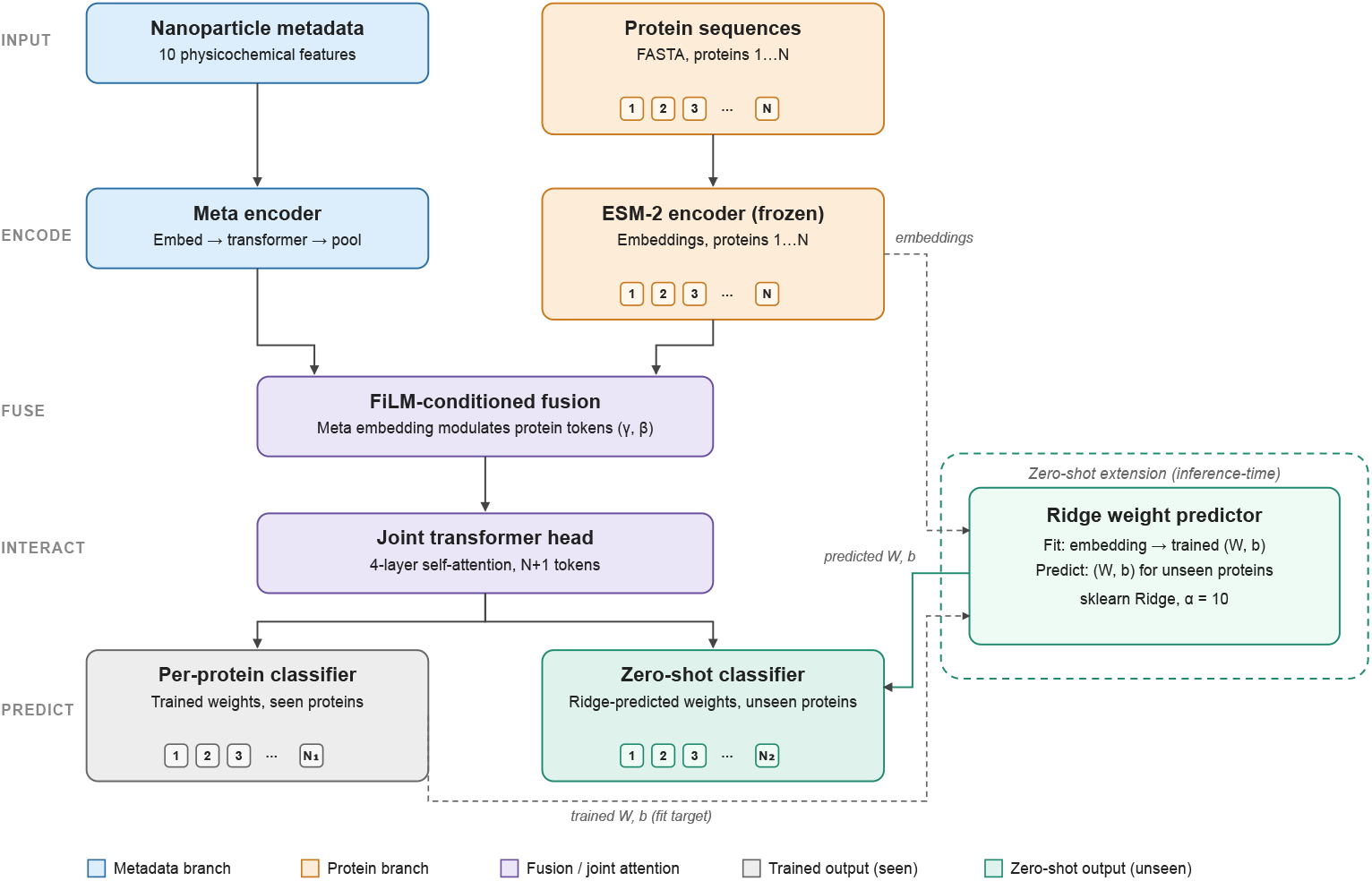
Overview of the NanoPCT framework. Nanoparticle metadata are encoded into a latent representation through a metadata encoder, while protein sequences are embedded by a frozen ESM-2 protein language model. A FiLM module conditions protein representations on nanoparticle properties before joint interaction is modeled by a transformer encoder. During training, NanoPCT learns protein-specific classifiers for all training proteins simultaneously. During inference, classifier parameters for unseen proteins are predicted from their ESM embeddings using ridge regression [35, 36], enabling zero-shot protein corona prediction without model retraining.

NanoPCT consists of two parallel encoding branches. The nanoparticle branch converts categorical and numerical physicochemical descriptors into a compact metadata embedding, whereas the protein branch extracts sequence representations using a frozen ESM-2 encoder [23]. Instead of directly concatenating these representations, NanoPCT employs Feature-wise Linear Modulation (FiLM) [37] to condition protein embeddings according to nanoparticle characteristics before introducing both modalities into a joint transformer. This design enables protein–protein interactions and nanoparticle-dependent modulation to be learned simultaneously within a shared latent space.

Unlike conventional multi-task learning frameworks, NanoPCT predicts adsorption probabilities for all proteins in a single forward pass. Furthermore, the learned classifier weights exhibit strong correspondence with protein sequence embeddings, allowing classifier parameters for unseen proteins to be reconstructed directly from sequence representations during inference. Consequently, NanoPCT naturally supports zero-shot prediction without requiring additional model optimization.

### 2.3 Understanding zero-shot generalization

#### 2.3.1 Protein sequence representation enables transferable prediction

A prerequisite for zero-shot prediction is that protein representations preserve biologically meaningful similarities while remaining compatible with supervised adsorption learning. To evaluate the contribution of sequence representations, we compared three protein encoding strategies: (i) frozen protein identity embeddings, (ii) trainable protein identity embeddings, and (iii) frozen ESM sequence embeddings.

As summarized in Table 1, replacing trainable protein identities with frozen ESM embeddings does not compromise predictive performance. Instead, ESM embeddings achieve the highest AUROC, AUPRC and F1-score among all evaluated representations.

**Table 1:** Comparison of different protein representations.

| Representation | OOF |  |  |
| --- | --- | --- | --- |
|  | AUROC | AUPRC | F1 |
| Protein ID (Frozen) | 0.9392 | 0.9369 | 0.8717 |
| Protein ID (Trainable) | 0.9401 | 0.9372 | 0.8720 |
| ESM embedding (Frozen) | **0.9405** | **0.9396** | **0.8743** |

Interestingly, trainable protein identity embeddings provide only marginal improvement over randomly initialized frozen identities, suggesting that protein identity alone is sufficient for fitting adsorption patterns of proteins observed during training. In contrast, pretrained sequence embeddings [23] retain equivalent predictive capability while encoding continuous biochemical relationships between proteins. This property enables NanoPCT to directly transfer adsorption knowledge from observed proteins to unseen proteins, thereby providing the representation basis for zero-shot prediction.

#### 2.3.2 Classifier weights are predictable in protein embedding space

NanoPCT performs zero-shot prediction by reconstructing classifier parameters for unseen proteins directly from their sequence embeddings. We therefore investigated whether classifier weights learned during training exhibit a continuous relationship with the underlying protein embedding space.

Figure 4 projects both learned classifier weights and ridge-reconstructed weights into the same low-dimensional space. Reconstructed classifiers closely overlap with the original classifiers throughout the embedding manifold, indicating that classifier parameters vary smoothly with protein sequence representations instead of forming isolated discrete clusters.

**Fig. 4:**
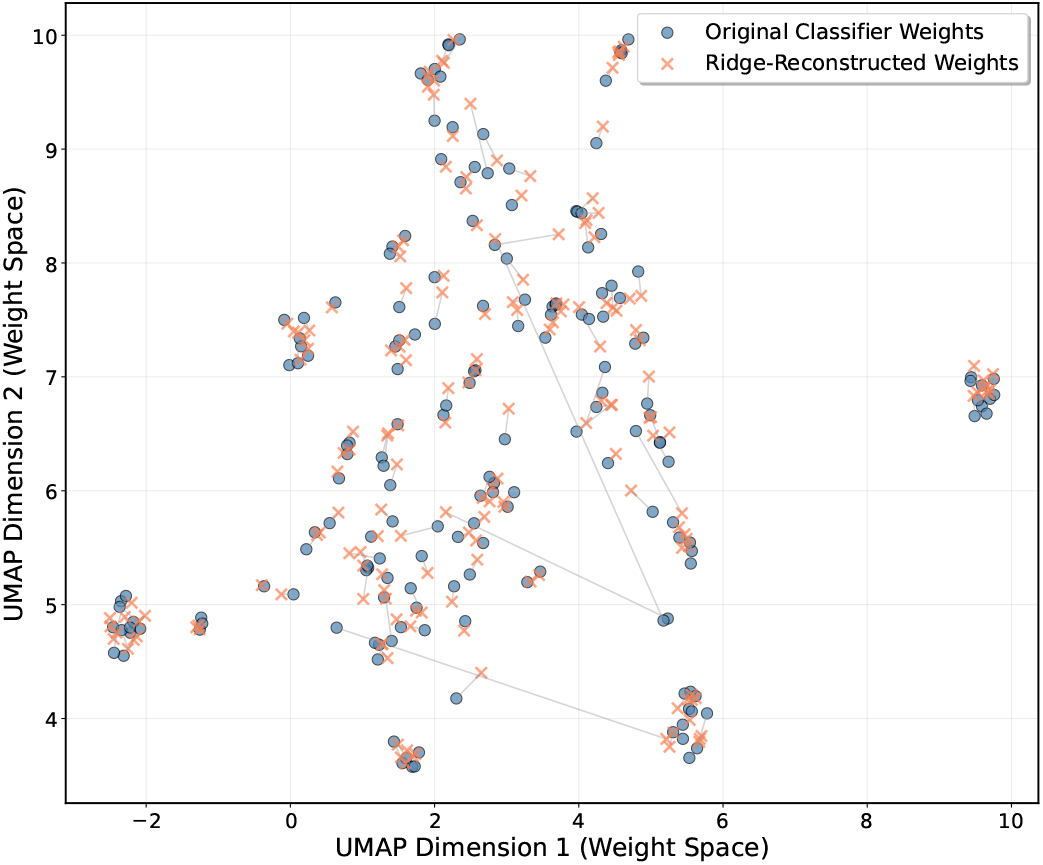
UMAP visualization of classifier weights. Blue circles denote classifiers learned during supervised training, while orange crosses represent classifier weights reconstructed from protein embeddings using ridge regression. Thin gray lines connect each original classifier weight to its corresponding reconstructed counterpart, illustrating the extent of deviation between the two sets of weights in the UMAP space.

To further evaluate different reconstruction strategies, we compared ridge regression with two alternative approaches, including nearest-neighbor averaging and global classifier averaging.

As shown in Table 2, reconstructing classifiers according to protein similarity substantially improves zero-shot performance compared with assigning identical classifier parameters to every unseen protein. While k-nearest-neighbor reconstruction already achieves competitive performance, ridge regression consistently produces the best overall accuracy.

**Table 2:** Comparison of zero-shot classifier reconstruction methods.

| Method | Zero-shot |  |
| --- | --- | --- |
|  | AUROC | AUPRC |
| Global averaging | 0.9040 | 0.8935 |
| kNN reconstruction | 0.9219 | 0.9164 |
| Ridge regression | **0.9222** | **0.9175** |

These observations suggest that protein sequence embeddings and classifier parameters share a common continuous geometry. Rather than memorizing individual proteins, NanoPCT learns a smooth mapping from protein sequence space to classifier space, enabling classifier parameters for previously unseen proteins to be inferred directly from their sequence representations.

#### 2.3.3 Functional-group-level zero-shot validation demonstrates biological transferability

To further evaluate whether NanoPCT can generalize beyond individual unseen proteins, we performed a more stringent functional-group-level zero-shot evaluation. Instead of randomly holding out proteins, all proteins belonging to the same biological functional group were completely excluded from training and used exclusively for validation. In each leave-one-functional-group-out fold, NanoPCT was trained using proteins from the remaining functional groups, while classifier parameters for the held-out proteins were reconstructed from their ESM sequence embeddings using ridge regression.

This evaluation represents a challenging distribution-shift scenario, where the model must transfer adsorption knowledge across biologically distinct protein classes rather than relying on interpolation among individual proteins with similar identities. As shown in Fig. 5, NanoPCT maintained strong predictive performance across most unseen functional groups. Across all leave-one-group-out evaluations, when averaging over all individual proteins, the model achieved an average AUROC of 0.8659, an average AUPRC of 0.8600, and an average F1-score of 0.7919.

**Fig. 5:**
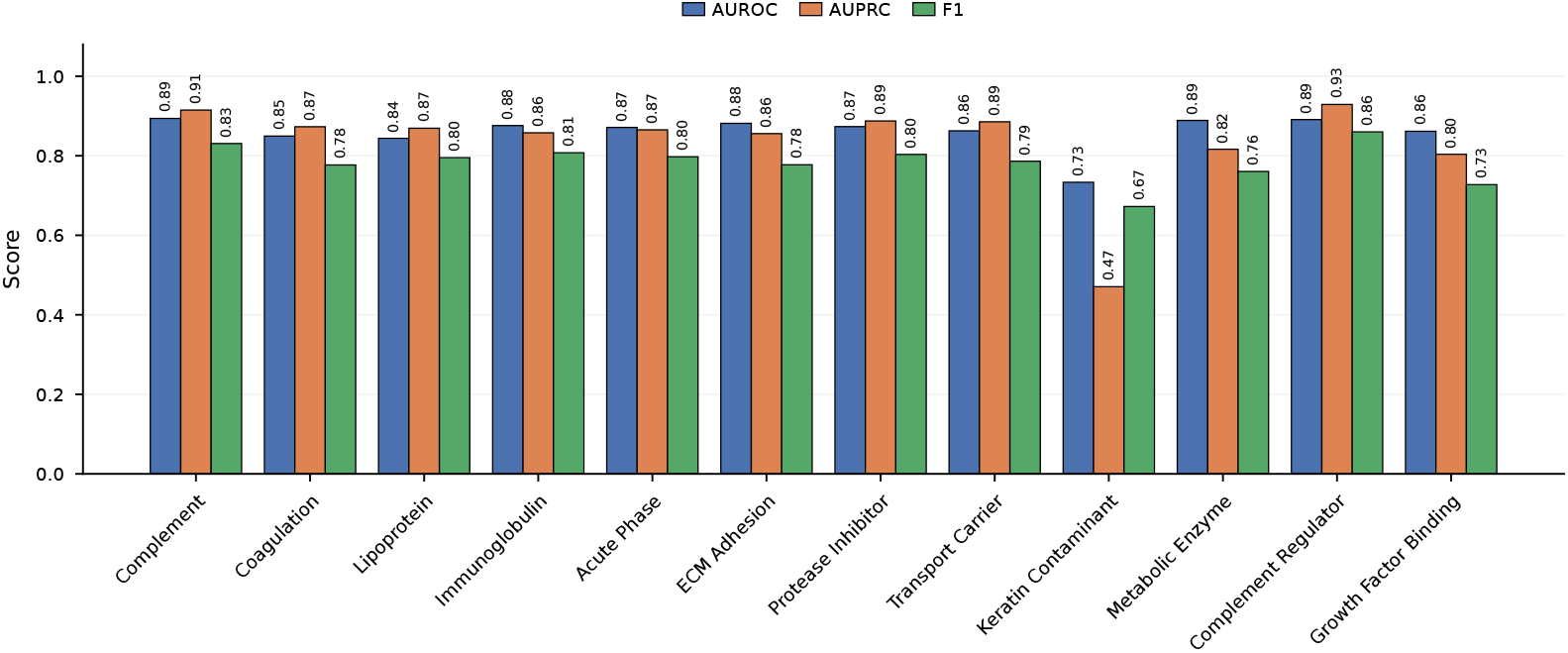
Functional-group-level zero-shot prediction performance. Each group was completely excluded from training and evaluated using ridge-based classifier reconstruction from protein sequence embeddings. AUROC, AUPRC and F1-score are shown for each held-out functional group.

Most functional groups exhibited consistently high prediction accuracy, with AUROC values ranging from 0.84 to 0.89 and AUPRC values generally exceeding 0.80. For example, Complement Regulator, Complement, Metabolic Enzyme, and Immunoglobulin groups achieved particularly strong transfer performance, demonstrating that adsorption patterns associated with these protein classes can be effectively inferred even when the corresponding functional groups are absent during training.

A notable exception was the Keratin Contaminant group, which showed reduced performance compared with other functional classes (AUROC = 0.73, AUPRC = 0.47, F1 = 0.67). This decreased accuracy likely reflects the distinct sequence characteristics and adsorption behavior of keratin-related proteins, which may occupy less represented regions of the learned protein embedding space.

Importantly, the overall preservation of prediction accuracy across diverse functional groups demonstrates that NanoPCT does not merely memorize protein-specific adsorption patterns. Instead, the model learns transferable relationships between protein sequence representations and nanoparticle adsorption behavior, enabling generalization to biologically distinct protein classes that were completely unseen during training.

Together with protein-level zero-shot experiments, these results provide stronger evidence that NanoPCT captures transferable relationships between protein sequence representations and nanoparticle adsorption behavior rather than protein-specific correlations.

### 2.4 Adaptive nano-protein interaction modeling improves protein corona prediction

To dissect the contribution of different interaction mechanisms to protein corona prediction, we performed an ablation study by evaluating four variants of the NanoPCT interaction module: (i) removing FiLM conditioning (w/o FiLM), (ii) replacing joint self-attention with cross-attention [34], (iii) replacing the interaction transformer with an independent MLP applied to each protein token, and (iv) removing all interaction modules and directly adding nanoparticle and protein representations. The quantitative comparison is summarized in Table 3.

**Table 3:** Ablation study of the interaction module. OOF evaluates prediction performance on proteins observed during training, while Zero-shot evaluates generalization to unseen proteins using Ridge-based classifier reconstruction.

| Model | OOF |  |  | Zero-shot |  |
| --- | --- | --- | --- | --- | --- |
|  | AUROC | AUPRC | F1 | AUROC | AUPRC |
| NanoPCT | 0.9405 | 0.9396 | **0.8743** | **0.9222** | **0.9175** |
| Cross-attention | **0.9420** | **0.9404** | 0.8735 | 0.9217 | 0.9163 |
| w/o FiLM | 0.9339 | 0.9335 | 0.8637 | 0.9159 | 0.9047 |
| Direct addition | 0.9298 | 0.9305 | 0.8621 | 0.9167 | 0.9098 |
| MLP replacement | 0.9129 | 0.9062 | 0.8410 | 0.9146 | 0.9059 |

Removing FiLM conditioning consistently reduced prediction performance, with the OOF AUROC decreasing from 0.9405 to 0.9339 and the zero-shot AUROC decreasing from 0.9222 to 0.9159. This indicates that nanoparticle-dependent modulation of protein representations plays an important role in capturing context-specific adsorption patterns. Unlike approaches that treat protein sequence embeddings as fixed descriptors, FiLM enables nanoparticle properties to dynamically regulate protein representations before interaction learning, thereby providing a more adaptive encoding of nano-bio interface states.

Replacing joint self-attention with cross-attention achieved comparable performance, with an OOF AUROC of 0.9420 and a zero-shot AUROC of 0.9217. This suggests that explicitly modeling dependencies between nanoparticle and protein representations is sufficient to capture strong conditional interactions. Nevertheless, the similar performance between cross-attention and NanoPCT indicates that joint self-attention provides an equally effective interaction mechanism while maintaining a unified representation space for simultaneously modeling multiple proteins associated with the same nanoparticle.

In contrast, directly adding nanoparticle and protein representations resulted in a notable performance decrease (AUROC = 0.9298). Although this operation preserves information from both modalities, it lacks the nonlinear transformation required to construct protein-specific adsorption states. The performance degradation suggests that protein corona formation is not determined by independent contributions from protein sequence and nanoparticle properties, but instead requires a learned interaction function that captures their conditional dependencies.

The MLP replacement exhibited the largest performance decline among all variants (AUROC = 0.9129, AUPRC = 0.9062), substantially underperforming both attention-based fusion strategies and direct addition. This result highlights that the effectiveness of the interaction module depends not only on nonlinear modeling capacity but also on the granularity and structure of feature integration. In this variant, each protein token is independently combined with the nanoparticle embedding and processed through an MLP pathway. Such independent fusion may obscure shared interaction patterns across proteins and introduce redundant feature combinations, making it more difficult for the model to identify biologically meaningful adsorption signals.

Collectively, these ablation results demonstrate that accurate protein corona prediction requires structured cross-modal interaction learning rather than simple feature aggregation. The consistent advantage of interaction-aware architectures supports the central design principle of NanoPCT: nanoparticle properties first provide contextual modulation of protein representations, followed by joint interaction learning to capture protein-specific adsorption behavior.

### 2.5 Inverse nanoparticle design reveals transferable adsorption principles

Having established the predictive accuracy and zero-shot generalization capabilities of NanoPCT, we next sought to investigate whether the model learns fundamental biophysical principles of nano-bio interactions that can be explicitly extracted and utilized for rational material design. To achieve this, we performed a large-scale *in silico* inverse design experiment [38, 39] targeting specific protein functional groups.

To systematically explore the parameter space, we employed Latin Hypercube Sampling (LHS) [40] to generate a highly diverse candidate pool of 100,000 nanoparticle design combinations, encompassing the full range of material features. For robust inference, we utilized a 5-fold ensemble of NanoPCT to predict the binding probability for each combination. To account for model uncertainty and prioritize reliable designs, we evaluated the binding capability using a conservative lower confidence bound score, defined as *µ* − 2*σ*, where *µ* and *σ* represent the mean and standard deviation across the 5-fold predictions, respectively.

We extracted the top 5% (optimal) and bottom 5% (suboptimal) combinations based on this conservative score for each protein functional group. To quantify the preference for specific categorical features (e.g., surface modification, shape, charge), we calculated the enrichment fold-change. Specifically, we computed log_2_(Top enrichment*/*Bottom enrichment), where a positive value indicates a strong selection bias for that feature category to achieve desired binding (**Fig. 6a, b**).

**Fig. 6:**
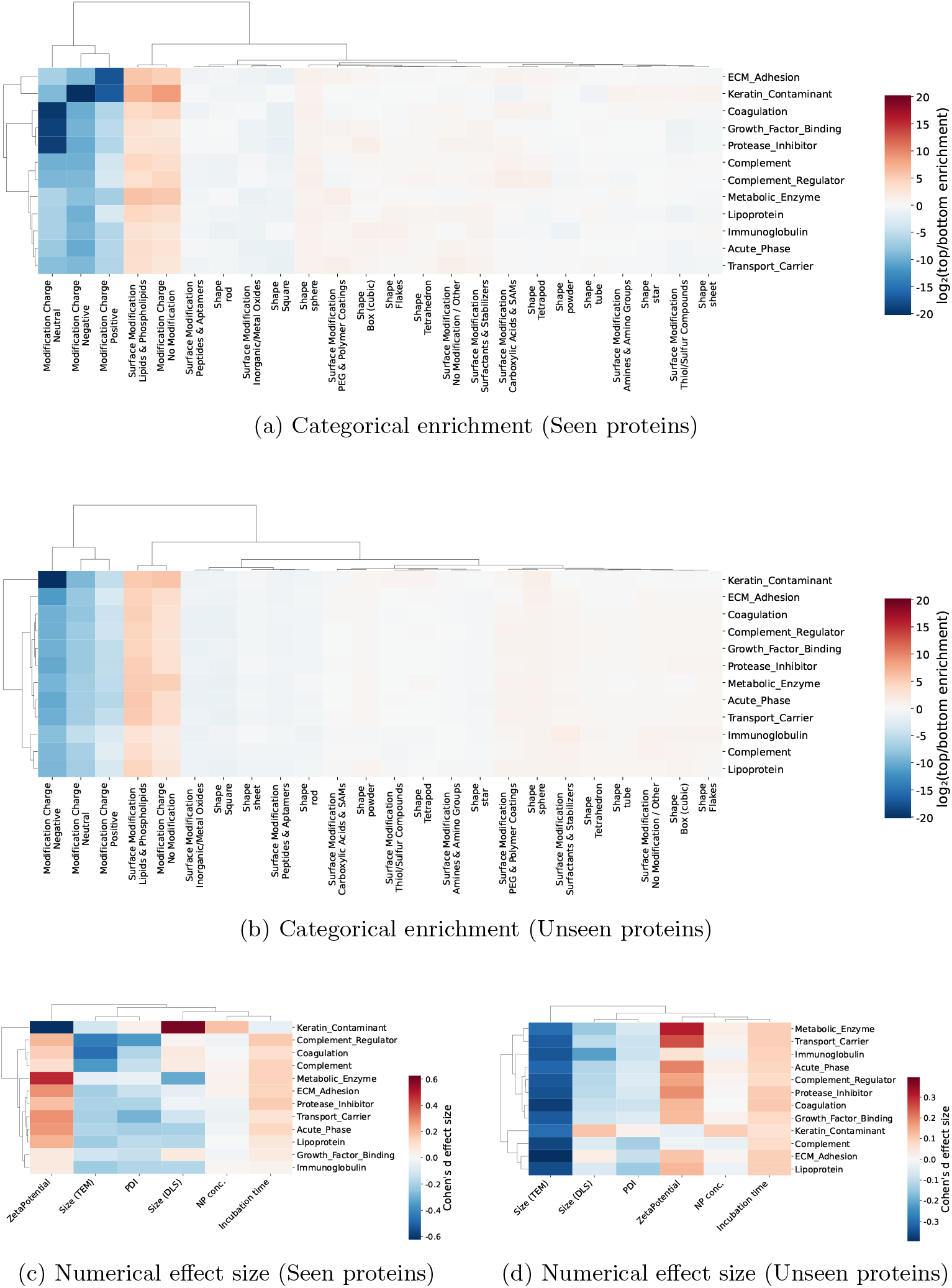
Transferable material design principles uncovered by large-scale *in silico* inverse sampling. **(a, b)** Hierarchical clustering heatmaps of log_2_(Top*/*Bottom) enrichment scores for categorical features (e.g., shape, surface modification) across distinct protein functional groups. A comparison between 178 trained (seen) proteins **(a)** and 215 zero-shot (unseen) proteins **(b)** demonstrates consistent combinatorial preferences. Red indicates positive enrichment in optimal designs, while blue indicates depletion. **(c, d)** Hierarchical clustering heatmaps of Cohen’s *d* effect sizes for numerical features (e.g., size, zeta potential) between top and bottom design conditions for seen **(c)** and unseen **(d)** proteins. Positive values (red) denote a preference for larger feature values, whereas negative values (blue) denote a preference for smaller values. The striking visual and statistical correlation between the left (seen) and right (unseen) panels validates the universality of the learned biophysical interactions.

For numerical features (e.g., size, zeta potential, incubation time), we assessed the design preference using Cohen’s *d* effect size [41] to eliminate scale disparities across different physical properties. The effect size was calculated as *d* = (*µ*_*top*_ − *µ*_*bottom*_)*/σ*_*pooled*_, where the pooled standard deviation is 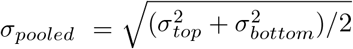. A large positive *d* value indicates that larger feature values are preferred for optimal binding, whereas a negative value suggests a preference for smaller values (**Fig. 6c, d**). Hierarchical clustering [42] of both categorical enrichments and numerical effect sizes revealed distinct, biologically coherent material design blueprints tailored to different protein functional classes.

Crucially, to demonstrate that these extracted design principles are universally applicable rather than merely overfitted to the training data, we rigorously compared the design landscapes between the 178 trained (“seen”) proteins and the 215 zero-shot (“unseen”) proteins. Strikingly, correlation analysis revealed a near-perfect consistency in the learned physical rules across the two sets. For numerical features, the mean consistencies achieved a Pearson *r* = 0.9994 and Spearman *ρ* = 1.0000 for the top conditions (72 effective paired groups), and *r* = 0.9818, *ρ* = 0.9770 for the bottom conditions. All correlations were highly significant (all *p <* 10^−15^).

Similarly, the categorical feature enrichment scores demonstrated robust transferability. The top enrichment consistency across 426 valid pairings yielded a Pearson *r* = 0.9844 and Spearman *ρ* = 0.9452, and the bottom enrichment consistency (432 valid pairings) yielded *r* = 0.8964, *ρ* = 0.8190, again with all *p*-values far below 10^−15^. This profound statistical concordance conclusively demonstrates that NanoPCT reveals transferable statistical principles underlying protein corona formation. Consequently, the generalizable design rules uncovered here can be reliably transferred to unseen proteins, offering a powerful predictive engine for the tailored engineering of nanomedicines.

## 3 Discussion

The predictive modeling of the nanoparticle–protein corona has long been constrained by the conventional “one-model-per-protein” paradigm [11, 16, 18–22]. In this formulation, each protein is treated as an independent prediction task, preventing the transfer of knowledge across proteins and making the prediction of previously uncharacterized proteins practically impossible. In this work, we introduce NanoPCT, which reconceptualizes protein corona prediction as a universal protein-conditioned learning problem. By directly integrating nanoparticle physicochemical properties with protein language model representations [23], NanoPCT learns a single transferable prediction function that not only performs robustly across diverse proteins but also generalizes to previously unseen proteins without retraining. This capability resembles zero-shot learning widely explored in computer vision [28, 43–45] and natural language processing [46], as well as at their intersection [47]. Our findings suggest that protein identity should not be regarded as the prediction target, but rather as part of the model input.

A key observation emerging from this study is that universal protein-conditioned learning is enabled by the continuous sequence representation learned by protein language models [23–25]. Our manifold analyses (Fig. 2a,b) show that unseen proteins and held-out functional groups are naturally represented within the same continuous sequence space occupied by training proteins rather than forming isolated regions. As the diversity of training proteins increases, feature-engineered machine learning models gradually lose predictive performance, whereas NanoPCT remains substantially more stable. Beyond individual protein generalization, functional-group-level zero-shot evaluation provides an even more stringent assessment of transferability. By excluding entire biological protein classes during training, NanoPCT still achieved robust prediction across most unseen functional groups. This observation indicates that the learned sequence representation and classifier reconstruction strategy capture shared biochemical determinants of nanoparticle adsorption that extend beyond local protein similarity. These findings suggest that pretrained sequence embeddings provide an effective biochemical manifold for sharing information across proteins [23], transforming what would otherwise be a difficult zero-shot extrapolation problem into a tractable local interpolation problem within the protein embedding space.

Beyond universal prediction, our results provide insight into why zero-shot prediction is possible. The ability to reconstruct classifier parameters directly from protein embeddings using a simple ridge regression model indicates that the learned decision boundaries vary smoothly across the protein embedding space. The successful prediction of completely unseen functional groups further supports this interpretation. If classifier reconstruction relied primarily on nearest-neighbor similarity between proteins, performance would be expected to substantially decrease when entire functional categories were removed. Instead, NanoPCT retained high predictive accuracy across diverse protein classes, suggesting that the embedding-to-classifier mapping reflects broader biological principles underlying protein–nanoparticle interactions. Proteins and protein groups occupying related regions of the ESM manifold tend to require similar classifier parameters, allowing adsorption decision functions learned from known proteins to be transferred to previously unseen ones. This observation suggests that pretrained protein representations capture biologically meaningful information that is directly relevant to nanoparticle–protein interactions and provide a practical foundation for parameter transfer without requiring additional model optimization [23].

Our ablation experiments further demonstrate that accurate protein corona prediction depends on learning context-dependent interactions between nanoparticles and proteins rather than simply combining their representations. Removing FiLM conditioning [37] or replacing the interaction module with simple additive fusion consistently reduced both in-distribution and zero-shot performance, indicating that nanoparticle properties must actively modulate protein representations before interaction modeling. Likewise, replacing the joint transformer with independent MLP processing substantially degraded performance, suggesting that explicit modeling of interactions across proteins is critical for learning shared adsorption patterns. Interestingly, replacing self-attention with cross-attention [34] produced comparable predictive accuracy, implying that both interaction mechanisms can effectively capture nanoparticle–protein relationships. Collectively, these results indicate that conditional feature modulation together with interaction-aware representation learning forms the computational foundation that enables both universal prediction and transferable zero-shot inference.

Perhaps the strongest evidence that NanoPCT captures transferable nano–bio interaction principles comes from the large-scale *in silico* inverse design experiments [38, 39]. By combining Latin Hypercube Sampling [40] with conservative lower-confidence-bound scoring, NanoPCT identified interpretable material design rules for different protein functional groups. Remarkably, both the categorical enrichment patterns and the numerical effect sizes inferred from the 178 training proteins were highly consistent with those obtained from the 215 completely unseen proteins, yielding near-identical design landscapes. This strong concordance suggests that the material preferences learned from observed proteins remain largely valid for previously unseen proteins. Consequently, NanoPCT is able to transfer design knowledge beyond the proteins encountered during training, providing a practical framework for rationally identifying nanoparticle properties that preferentially recruit or avoid specific classes of proteins.

Despite these encouraging results, several limitations remain. First, although sequence-based protein language models provide a powerful representation of biochemical properties, protein corona formation is ultimately governed by three-dimensional molecular structures and surface accessibility [48–50]. Integrating structure-aware representations derived from geometric deep learning [51] or AlphaFold [48, 49] may further improve the modeling of conformationally dependent interactions. Second, the current framework predicts adsorption for individual proteins independently, whereas physiological coronas arise through dynamic competitive adsorption among many proteins, such as those described by the Vroman effect [52]. Extending NanoPCT toward multi-protein competitive prediction therefore represents an important direction for future research. Finally, important biological factors, including post-translational modifications, serum composition, ionic strength, and other environmental conditions [8], are not explicitly modeled in the current framework and may influence corona formation under realistic biological environments.

In conclusion, NanoPCT establishes a generalizable and interpretable computational framework for protein-conditioned nanoparticle corona prediction. More broadly, our findings suggest that protein language models provide a practical foundation for unifying molecular biology and nanomaterial informatics within a single transferable predictive framework. By combining universal prediction, zero-shot generalization across diverse protein classes, and interpretable inverse design, NanoPCT shifts protein corona modeling from empirical protein-specific predictors toward a sequence-aware computational paradigm capable of supporting the rational design of next-generation nanomaterials.

## 4 Methods

### 4.1 Datasets and data preprocessing

#### Data curation

To construct a comprehensive and highly diverse nanoparticle-protein corona dataset, we integrated data from 91 peer-reviewed publications spanning the years 2000 to 2024, primarily consolidating established databases [11, 16, 21, 22], alongside newly curated records from 2025. The initial aggregated dataset cataloged the relative protein abundance (RPA) of over 800 distinct proteins. However, the majority of these proteins were reported in only a single experimental study, lacking the statistical commonality required for robust machine learning. To ensure data quality, we applied a strict missing-value filter, excluding proteins with a sparsity (missing rate) exceeding 96% across all nanoparticle conditions. This filtering process yielded a refined matrix of 671 unique nanoparticle experimental conditions and 397 quantifiable proteins.

#### Feature engineering and label binarization

Each nanoparticle condition is described by a set of physicochemical descriptors, which were explicitly divided into categorical and numerical features. Categorical features include *Subtype, Surface Modification, Modification Charge*, and *Shape*. Numerical features encompass *Size (TEM), Size (DLS), Zeta Potential, Incubation NP concentration, Incubation time*, and *Polydispersity Index (PDI)*. Missing numerical values were imputed with a placeholder −1) prior to model ingestion. Following the established protocol by Ban et al. [16], we formulated the protein corona prediction as a binary classification task. Specifically, proteins with an RPA *>* 0 were assigned a positive label (1, denoting binding), whereas an RPA = 0 was assigned a negative label (0, denoting no binding). This binary formulation effectively mitigates quantification discrepancies across disparate proteomic measurement techniques (e.g., variants of LC-MS/MS or gel electrophoresis [12, 13]) sourced from different studies. Missing labels for specific proteins under certain experimental conditions were dynamically masked out during loss calculation.

#### Data splitting for universal and zero-shot learning

To evaluate both the universal learning capacity and the zero-shot generalization of NanoPCT, the 397 retained proteins were meticulously partitioned based on their data abundance:

- **Seen proteins (Training set):** 178 proteins with highly enriched data (missing rate *<* 23%), yielding a total of 95,429 valid nanoparticle-protein interaction pairs, were utilized for supervised universal training.
- **Unseen proteins (Zero-shot test set):** 215 proteins with higher sparsity (23% *<* missing rate *<* 96%), yielding 8,402 valid zero-shot interaction pairs, were strictly isolated from the training phase. These proteins were reserved exclusively to evaluate the model’s zero-shot predictive capability.
- **Excluded proteins:** 4 proteins satisfying the missing rate criteria were ultimately discarded because they exhibited single-class labels (e.g., positive in all available samples). Such lack of variance precludes the calculation of discriminative metrics such as AUROC and AUPRC.

Consequently, the final dataset comprises 178 training proteins and 215 zero-shot testing proteins.

### 4.2 Problem formulation for universal protein-conditioned learning

Traditional computational approaches for predicting the nanoparticle-protein corona rely on training independent models for individual proteins [11, 16, 18–22]. Formally, given a dataset of nanoparticle physicochemical features *X*, prior methods learn a distinct mapping function *f*_*i*_ : *X*→ *Y*_*i*_ for the *i*-th protein. In this formulation, an independent prediction function is trained for each protein.

We formulated protein corona prediction as a universal protein-conditioned learning problem. Instead of multiple isolated models, we aimed to learn a single, unified mapping function *F* : (*X, P*) →*Y*, where *X* represents the nanoparticle properties and *P* represents the intrinsic characteristics of a given protein.

To validate the feasibility of this universal formulation before deploying deep learning architectures, we established robust machine learning baselines (e.g., LightGBM [30] and XGBoost [31]). For these tree-based baselines, the protein characteristics *P* were explicitly featurized using a comprehensive set of 431 descriptors, following the approach of Huzar et al.[19] (detailed in Supplementary Table S1). Protein descriptors comprised global physicochemical properties (e.g., molecular weight, isoelectric point, GRAVY), structural fractions, flexibility indices, as well as multiscale compositional representations (including 20 single amino acid and 400 dipeptide frequencies). The nanoparticle features and the 431-dimensional protein descriptor vectors were concatenated to form a flattened input vector. To assess capacity limits and negative transfer, these baseline models were subjected to an escalating curriculum of target proteins (random subsets of 20, 40, 80, 120, and all 178 training proteins) using a strict 10-fold cross-validation protocol.

### 4.3 NanoPCT architecture

NanoPCT consists of three core components: the nanoparticle MetaEncoder, the protein sequence language model, and a joint interaction head guided by Feature-wise Linear Modulation (FiLM).

#### 4.3.1 Nanoparticle feature encoding via tokenized transformers

Nanoparticle features were encoded using a token-based MetaEncoder adapted from FT-Transformer [53].

##### Categorical and numerical tokenization

For categorical descriptors, standard embedding lookup tables were utilized to project each discrete category into a dense vector of dimension *D*. For numerical descriptors, we adopted a *Fourier Encoding* mechanism proposed by Tancik et al. [54]. Specifically, each raw numerical scalar *x* is projected into a multi-frequency continuous space using *F* = 4 frequency bands:

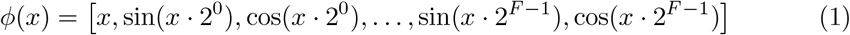

The resulting (2*F* + 1)-dimensional Fourier representation is then mapped through a linear layer and Layer Normalization [55] to match the embedding dimension *D*.

##### Feature interaction and attention pooling

The embedded categorical and numerical tokens are concatenated along the feature sequence dimension to form a nanoparticle token matrix *X* ∈ ℝ^*N* ×*D*^, where *N* is the total number of features. This matrix is processed by a stack of standard Transformer blocks [34], utilizing Multi-Head Self-Attention (MHSA) and Gaussian Error Linear Unit (GELU) activations [56].

Finally, we designed a *Feature Attention Pooling* module [57] to adaptively aggregate the token sequence into a single, fixed-length nanoparticle representation vector. A learned attention network assigns a weight *α*_*i*_ to each feature token:

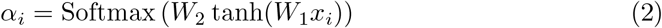

The final nanoparticle latent representation is the weighted sum of all feature tokens, 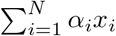, which is subsequently passed through a projection MLP to match the hidden dimension required for protein condition fusion.

#### 4.3.2 Protein sequence embedding via Pre-trained Language Models

Protein sequences were embedded using the pretrained ESM-2 model [23]. The 35M parameter model was used because it provided a balance between embedding dimensionality and computational cost.

ESM parameters remained frozen throughout training. To derive a fixed-length representation for the entire protein, we applied masked mean pooling over the sequence dimension:

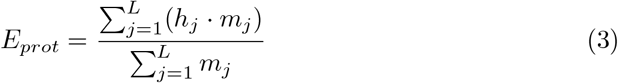

where 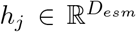 is the hidden state of the *j*-th amino acid token from the final ESM-2 layer, *m*_*j*_ ∈ {0, 1} is the attention mask, and *D*_*esm*_ = 480 is the embedding dimension.

#### 4.3.3 Feature-wise Linear Modulation (FiLM) and Joint Interaction Transformer

Protein embeddings were conditioned using Feature-wise Linear Modulation (FiLM) [37].

Let *E*_*meta*_ ∈ R^*D*^ denote the pooled nanoparticle representation generated by the MetaEncoder, and 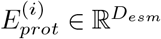 denote the sequence embedding of the *i*-th protein. The FiLM layer linearly projects *E*_*meta*_ to generate scale (*γ*) and shift (*β*) vectors, which conditionally modulate the protein embeddings:

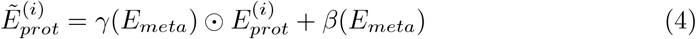

where ⊙ denotes element-wise multiplication. To align dimensions, *E*_*meta*_ is mapped to *D*_*esm*_ prior to modulation.

Following FiLM conditioning, a *Joint Transformer Head* [34] is deployed to model the complex, high-order interactions. The modulated protein embeddings for all *N* target proteins are concatenated with the linearly projected nanoparticle token, forming a combined sequence tensor 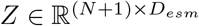. This tensor is processed by a stack of *L* = 4 standard Transformer encoder layers, utilizing 16 attention heads.

#### Protein-specific linear classification

Upon exiting the joint transformer layers, the updated protein tokens 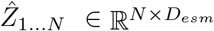 (discarding the first token representing the material) are passed to a linear classification layer. Instead of using a shared MLP across all proteins, we defined a trainable weight matrix 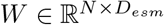 and bias vector *b* ∈ ℝ^*N*^. The final adsorption logits for the *N* proteins are computed via Einstein summation:

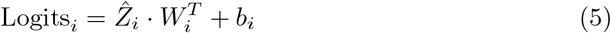

This design ensures that each training protein *i* learns an independent, continuous classifier weight vector *W*_*i*_.

### 4.4 Ablation study configurations

To systematically dissect the contribution of each architectural innovation in NanoPCT, we designed and trained several ablated variants (detailed in Supplementary Note S2). For rigorous comparability, all variants seamlessly interface with the *N* protein-specific linear classifiers to support the identical zero-shot evaluation pipeline. The key ablations included:

- **Alternative protein representation strategies:** Replacing the frozen ESM-2 embeddings with naive, non-sequential identity-based representations (Frozen and Trainable Protein ID Embeddings) to validate the necessity of language model semantics.
- **Joint interaction module ablations:** Systematically altering the fusion mechanism by (1) removing the FiLM conditioning, (2) substituting the standard Joint Transformer with a Cross-Attention module, (3) reducing fusion to a direct element-wise addition, or (4) replacing the interaction transformer entirely with a standard Multilayer Perceptron (MLP) mapping.

These variants were evaluated under the exact same 10-fold cross-validation protocols as the full NanoPCT model.

### 4.5 Model training and zero-shot inference

#### 4.5.1 Supervised training on seen proteins

NanoPCT was initially trained exclusively on the 178 data-rich proteins. The training process utilized a strict 10-fold cross-validation scheme to ensure robust evaluation and prevent overfitting. Because not all proteins were measured in every nanoparticle experiment, we employed a masked Binary Cross-Entropy (BCE) loss:

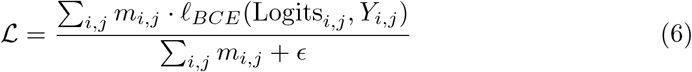

where *Y*_*i,j*_ is the true binding label of the *j*-th protein under the *i*-th nanoparticle condition, *m*_*i,j*_ ∈ {0, 1} is a binary mask indicating whether this specific measurement exists in the dataset, and *ϵ* (e.g., 1 × 10^−8^) is a small constant to prevent division by zero. After cross-validation, the trained weights from all 10 folds were saved independently for zero-shot inference.

#### 4.5.2 Zero-shot generalization via classifier reconstruction

During supervised training, the model learns a task-specific classifier weight matrix 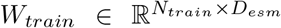 and bias vector 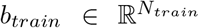 corresponding to the predefined sequence coordinate space 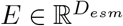.

We evaluated three mathematical strategies to reconstruct the classifiers (*W*_*unseen*_, *b*_*unseen*_) for the *N*_*unseen*_ = 215 zero-shot proteins:

1. **Global Averaging:** Assuming no sequence-specific dependency, assigning the average weights of all training proteins to novel proteins:

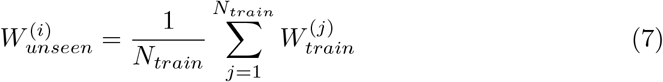
2. *K***-Nearest Neighbors:** Employing distance-weighted averaging of the *k* = 15 closest neighbors in the ESM-2 space based on cosine similarity:

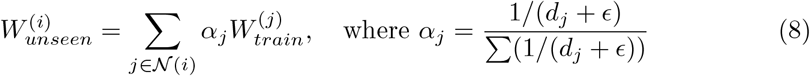
3. **Ridge Regression [36]:** Capturing the global continuous transformation via *L*_2_-regularized linear regression:

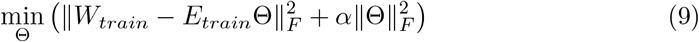

where ∥ · ∥_*F*_ denotes the Frobenius norm, 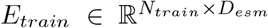 is the training sequence embedding matrix, and 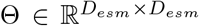 is the transformation matrix to be learned. Once fitted (*α* = 10), weights for unseen proteins were predicted via *W*_*unseen*_ = *E*_*unseen*_Θ.

During zero-shot inference, the original classification layer was procedurally replaced with the reconstructed parameters. The nanoparticle features and the frozen unseen protein embeddings were passed forward through the network across all 10 folds without retraining.

#### 4.5.3 Functional-group-level zero-shot evaluation

To further evaluate the biological transferability of NanoPCT under a more challenging distribution-shift scenario, we performed a leave-one-functional-group-out zero-shot evaluation. Unlike conventional protein-level zero-shot prediction, where individual unseen proteins are excluded from training, this protocol removes all proteins belonging to an entire functional group during model training.

Functional groups were assigned according to UniProt functional annotations in the nanoparticle-protein corona dataset. For each evaluation fold, one functional group was selected as the held-out group, and all proteins belonging to this group were completely excluded from supervised training. The remaining functional groups were used to train NanoPCT following the same supervised learning procedure described above.

After training, classifier parameters for proteins within the held-out functional group were reconstructed without additional optimization. Specifically, the learned relationship between ESM-2 protein embeddings and protein-specific classifier parameters was used to predict the corresponding classifier weights and biases for the unseen proteins. Ridge regression was employed as the classifier reconstruction strategy:

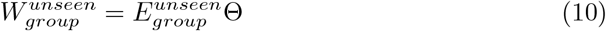

where Θ represents the embedding-to-classifier transformation matrix learned exclusively from proteins belonging to the remaining functional groups.

The reconstructed classifiers were then inserted into the frozen NanoPCT model, and predictions were generated for the held-out functional group without any parameter updating or exposure to its interaction labels. This procedure was repeated iteratively until every functional group had served as the unseen test group.

Performance was evaluated using AUROC, AUPRC, and F1-score following the same macro-averaging strategy described in the evaluation metrics section. The final functional-group-level zero-shot performance was reported as the average across all leave-one-group-out evaluations.

### 4.6 *In silico* inverse design and statistical analysis

#### Generation of candidate design space via Latin Hypercube Sampling

Candidate nanoparticle designs were generated using Latin Hypercube Sampling [40]. **Ensemble inference and robust conservative scoring**. Prediction probabilities were aggregated across the five folds with the highest zero-shot performance. For each candidate and target protein pair, we calculated the mean prediction probability (*µ*) and the standard deviation (*σ*) across the 5 folds. We defined a conservative Lower Confidence Bound (LCB) score [58] as *µ −* 2*σ*. For inverse design [38, 39] targeting zero-shot proteins, the necessary classifier weights were dynamically reconstructed via the aforementioned ridge regression mapping prior to executing the 5-fold forward pass.

#### Statistical enrichment and effect size analysis

For categorical features, enrichment fold-change was calculated as log_2_(Top 5% Frequency*/*Bottom 5% Frequency) to quantify the selection bias toward specific categorical levels within the theoretically optimal versus suboptimal designs. For continuous features, Cohen’s *d* effect size (*d* = (*µ*_*top*_ *− µ*_*bottom*_)*/σ*_*pooled*_) [41] was utilized. Hierarchical clustering (Ward’s method, Euclidean distance) [42, 59] was performed on enrichment scores and effect sizes.

### 4.7 Implementation details and evaluation metrics

#### Evaluation Metrics and Thresholding

Model performance on training proteins was evaluated using 10-fold out-of-fold (OOF) prediction. Zero-shot performance was evaluated independently for each trained fold using fold-specific classifier reconstruction, and the reported results correspond to the average across all folds.

Given that nanoparticle-protein datasets suffer from sparsity and class imbalance, Area Under the Precision-Recall Curve (AUPRC) was prioritized alongside the Area Under the Receiver Operating Characteristic curve (AUROC) and F1-score. All reported metrics were calculated using a *macro-averaging* strategy: metrics were computed independently for each individual protein before calculating the unweighted arithmetic mean. For the F1-score, an optimal decision threshold *t* ∈ [0.05, 0.95] was dynamically determined via grid search to maximize the score while satisfying a strict lower bound on Recall (≥ 0.6).

#### Training optimization and environment

The NanoPCT model and its variants were trained end-to-end using the AdamW optimizer [60] coupled with a `SequentialLR` scheduler (linear warmup followed by Cosine Annealing decay [61]). To counteract the inherent label imbalance, a specific positive weight (*w*_*pos*_) was dynamically calculated based on the negative-to-positive sample ratio and integrated into the masked loss function. Comprehensive hyperparameter configurations, including learning rates, dropout ratios, batch sizes, and gradient clipping thresholds, are detailed in Supplementary Note S3. All experiments were executed on a single NVIDIA GeForce RTX 5090 GPU (32 GB VRAM). The computational pipeline was implemented in Python (v3.10) using PyTorch (v2.8.0) and Automatic Mixed Precision (AMP) [62] to accelerate execution.

## 4.8 Data and code availability

The curated nanoparticle-protein corona dataset, the source code of the NanoPCT framework will be made publicly available on GitHub at https://github.com/issair/NanoPCT.

## Supplementary

### S1: Summary of protein descriptors

**Table S1:**
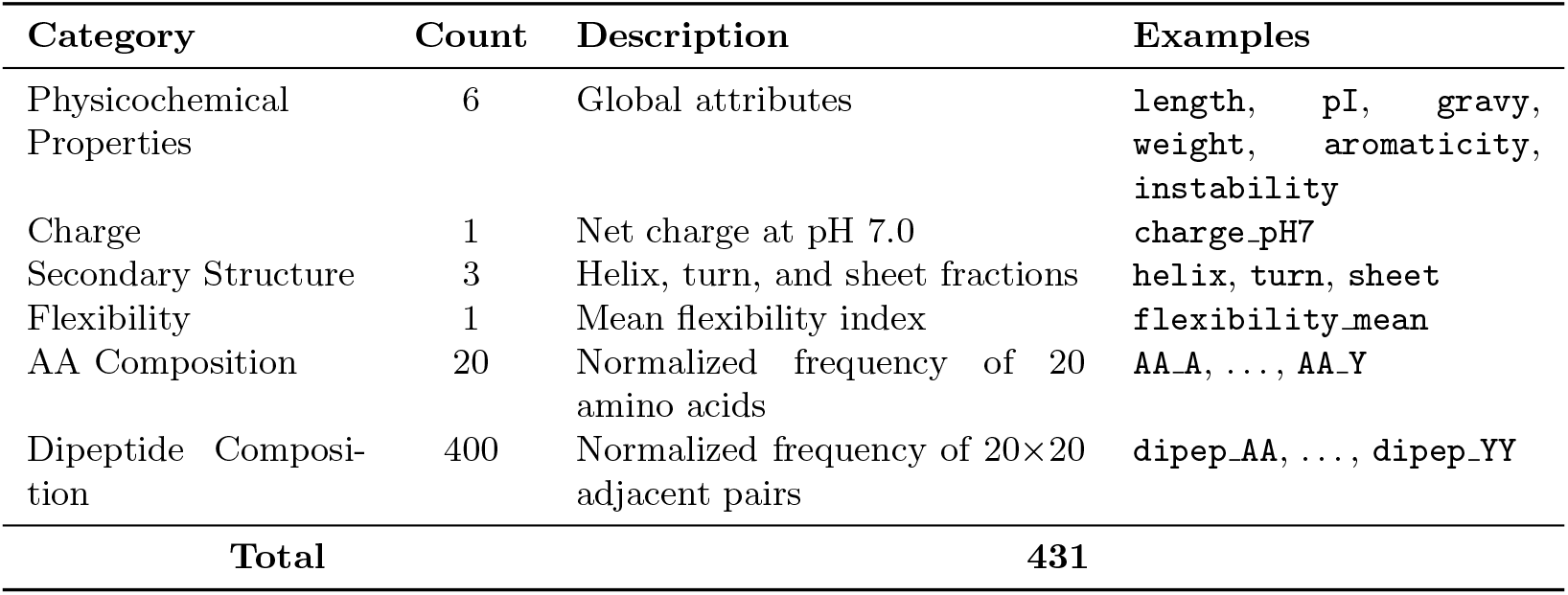
Summary of protein descriptors.

### S2: Ablation study configurations

To systematically dissect the contribution of each architectural innovation and representation strategy in the NanoPCT framework, we designed and trained several ablated variants. For rigorous comparability, all variants were structurally constrained to output exactly *N* updated protein tokens of dimension *D*_*esm*_, ensuring they seamlessly interface with the subsequent *N* protein-specific linear classifiers and support the identical zero-shot evaluation pipeline.

### Alternative protein representation strategies

To validate whether the model genuinely benefits from the biochemical semantics encoded by the ESM-2 [23] language model—rather than simply memorizing discrete protein identities from the training distribution—we compared the frozen ESM-2 embeddings against two non-sequential, identity-based baseline representations:

- **Frozen Protein ID Embedding:** Each of the *N* training proteins was assigned a unique, randomly initialized embedding vector sampled from a standard normal distribution *N* (0, 0.02). These embeddings were strictly frozen during training, serving as a naive baseline entirely devoid of sequence, structural, or evolutionary context.
- **Trainable Protein ID Embedding [29]:** Identical initialization to the frozen ID embeddings, but the vectors were actively updated during backpropagation. This configuration represents the conventional multi-task learning approach.

All other network components (FiLM, Joint Transformer [34]) remained identical to the standard NanoPCT. It should be noted that while ID-based embeddings can be effectively evaluated for out-of-fold (OOF) performance on seen proteins, they intrinsically lack a continuous sequence manifold, rendering the mathematical reconstruction of zero-shot parameters (via Ridge Regression or KNN) ill-posed.

### Joint interaction module ablations

To isolate the effects of feature modulation and self-attention mechanisms on capturing nano-bio physical interactions, we designed the following architectural variations for the interaction head:

- **Without FiLM conditioning [37]:** The Feature-wise Linear Modulation (FiLM) layer was removed. The raw, unmodulated protein ESM embeddings (*E*_*prot*_) were directly concatenated with the nanoparticle meta-token (*E*_*meta*_) to form the (*N* +1) sequence tensor, which was then fed into the standard Joint Transformer.
- **Cross-attention fusion [34]:** The standard self-attention Joint Transformer was replaced with a Cross-Attention module (4 layers, 16 attention heads). In this configuration, the unmodulated protein tokens 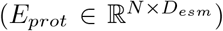 served strictly as the *Queries*, while the nanoparticle meta-token 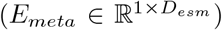 served as the *Keys* and *Values*.
- **Direct addition:** Both FiLM and the attention-based transformers were completely removed. The fusion process was reduced to a simple, naive element-wise addition broadcasted across the protein dimension: 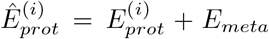. The resulting *N* fused tokens were passed directly to the linear classifiers.
- **MLP replacement:** The Joint Transformer was substituted with a standard Multilayer Perceptron (MLP). To maintain the *N* distinct protein pathways, the meta-token *E*_*meta*_ was concatenated with each modulated protein token individually. Each concatenated vector 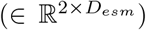 was processed independently by a 3-layer MLP with hidden dimensions of [1024, 512, 256], and finally projected back to dimension *D*_*esm*_ to feed into the respective classifiers.

### S3: Training optimization and environment

To ensure the reproducibility of our universal protein-conditioned learning framework, we detail the comprehensive hyperparameter configurations, regularization strategies, and optimization schedules utilized during the training of the NanoPCT model.

#### Optimization and learning rate scheduling

The models were trained end-to-end using the AdamW optimizer [60], which decouples weight decay from the gradient update to provide better generalization. The optimization hyperparameters were configured as follows:

- **Base learning rate:** 5 × 10^−4^
- **Weight decay:** 1 × 10^−4^
- **Momentum parameters:** *β*_1_ = 0.9, *β*_2_ = 0.999

To stabilize the optimization trajectory, particularly during the initial feature alignment phase between the untrained MetaEncoder and the frozen ESM-2 embeddings [23], we implemented a `SequentialLR` scheduler over a total training duration of 20 epochs. The learning rate was linearly warmed up [61] from 1 × 10^−6^ to the base learning rate (5 × 10^−4^) over the first 5 epochs. Subsequently, a Cosine Annealing learning rate decay was applied from epoch 6 to epoch 20, gradually reducing the learning rate down to a minimum threshold of 2 × 10^−4^.

#### Batching and regularization strategies

Due to the memory constraints imposed by modeling *N* = 178 simultaneous protein interactions per material input, we employed gradient accumulation to simulate a larger batch size.

- **Effective batch size:** A physical batch size of 16 was utilized, combined with 2 gradient accumulation steps, yielding an effective batch size of 32 for each optimization step.
- **Gradient clipping [63]:** To prevent exploding gradients during the highly parameterized transformer updates, global gradient clipping was applied with a maximum *L*_2_ norm threshold of 1.0.
- **Dropout [64]:** To mitigate overfitting given the limited nanoparticle sample size, dropout layers with a probability of *p* = 0.1 were systematically applied within the MetaEncoder sequence processing, the FiLM conditioning pathways, and the Joint Transformer modules.

#### Loss function and class imbalance handling

The dataset exhibits significant label imbalance, as protein non-binding events vastly outnumber binding events across the experimental conditions. To prevent the model from converging to a trivial majority-class predictor, a class-balanced masked Binary Cross-Entropy (BCE) loss was utilized. For each specific training protein *i*, a positive weight scalar 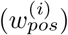 was independently calculated based on its training set statistics:

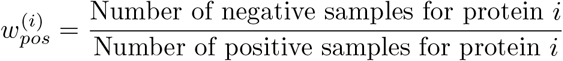

This 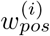 was passed to the `pos_weight` argument of the PyTorch `BCEWithLogitsLoss`function, ensuring that rare binding events received proportionally higher penalty gradients during backpropagation.

